# TAF4b Transcriptional Regulation During Cellular Quiescence of Developing Prospermatogonia

**DOI:** 10.1101/2022.04.20.488869

**Authors:** Megan A. Gura, Myles A. Bartholomew, Kimberly M. Abt, Soňa Relovská, Kimberly A. Seymour, Richard N. Freiman

## Abstract

Prospermatogonia (ProSpg) link the embryonic development of male primordial germ cells to the healthy establishment of postnatal spermatogonia and long-term mammalian spermatogenesis. While these spermatogenic precursor cells undergo the characteristic transitions of cycling and quiescence, the transcriptional events underlying these developmental hallmarks remain unknown. Here we investigated the expression and function of TAF4b in the timely development of mouse ProSpg using an integration of gene expression profiling and chromatin mapping. We find that *Taf4b* mRNA expression is elevated during the transition of M-to-T1 ProSpg and *Taf4b-*deficient ProSpg are delayed in their entry into quiescence. Gene ontology, protein network analysis, and chromatin mapping demonstrate that TAF4b is both a direct and indirect regulator of cell cycle-related gene expression programs during ProSpg quiescence. By comparing the transcriptome changes in male and female *Taf4b-*deficient embryonic germ cells, we revealed that TAF4b promotes sex-independent and -dependent gene expression pathways, highlighting its unique and critical role in the fertility of both sexes.

## INTRODUCTION

Male fertility is dependent on a highly regulated series of developmental events that begin with primordial germ cell (PGC) specification in early fetal development and end with the exhaustion of an adult unipotent spermatogonial stem cell (SSC) population during old age [1–3]. In mammals, PGC specification is achieved via BMP signaling to a small group of dorsally localized extra-embryonic cells; this BMP signaling then induces the expression of the master transcription regulators PRDM1, PRDM14, and TFAP2C [4–6]. While more heterogeneous in nature, several transcription factors mark and/or support adult SSC identity, including ID4, RHOX10, PAX7, and SALL4 [7–13]. Prospermatogonia (ProSpg, also called ‘gonocytes’) are male germ cells that have differentiated past PGC specification and have the potential to differentiate into adult spermatogonia (Spg) or become part of the SSC pool which provides the long-term renewing capabilities of the testis. Several critical molecular events occur during ProSpg development. First, epigenetic marks that were erased in PGCs are re-established during ProSpg development. Second, an initial pool of unipotent SSCs is thought to arise from the heterogeneous pool of developing ProSpg during this developmental window. These events are necessary for the correct establishment of spermatogenesis and its maintenance throughout adulthood, but the underlying gene expression networks that correctly time and integrate these complex events are currently unknown.

The expression of the transcription factors OCT4, KLF4, SOX2, and NANOG, which orchestrate the reprogramming of differentiated cells into induced pluripotent stem cells (iPSCs), are an excellent example of how the regulation of RNA polymerase II (Pol II) transcription is critical for many developmental transitions [14–16]. These sequence-specific DNA-binding proteins are thought to recruit Pol II to specific core promoters via the global transcription machinery and this transcriptional reprogramming leads to resetting the epigenome to a pluripotent state. How the regulation of Pol II transcription is achieved in ProSpg and how it is integrated with or instructive for epigenetic reprogramming of the male germline genome at these specific stages of development is unknown. We have identified a unique germ cell-enriched subunit of the general transcription factor TFIID, called TAF4b, that is required for proper long-term spermatogenesis in the mouse. Male mice lacking TAF4b become infertile after undergoing an SSC-independent first round of spermatogenesis; however, spermatogenesis can be re-established in the *Taf4b*-deficient testis following transplant of wildtype SSCs reflecting a functional adult SSC niche [17]. More recent characterization of spermatogenesis in our *Taf4b*-deficient mice revealed that TAF4b is required for embryonic male germ cell development and helps regulate the delicate balance of SSC self-renewal and differentiation [17,18]. In addition to our studies of TAF4b in mice, men who express a truncated version of the TAF4b protein are infertile owing to progressive azoospermia [19], and single nucleotide polymorphisms in human TAF4b have been linked to nonobstructive azoospermia [20]. Thus, deciphering the developmental and molecular mechanisms of TAF4b in the mouse can improve our understanding and model important aspects of male reproductive development and fertility in boys and men.

To contextualize the phenotype of *Taf4b*-deficient male germ cells, we examined the spatial and temporal expression of TAF4b during embryonic testis development. We analyzed previously published RNA-seq data sets derived from *Oct4*-eGFP mice, in which germ cells were separated from somatic cell populations through fluorescence-activated cell sorting (FACS; [21]). We determined that levels of *Taf4b* mRNA become progressively and significantly elevated in a germ cell-specific manner from embryonic day (E) 11.5 to E18.5. TAF4b protein expression was germ cell-specific at E13.5 and suggests it plays an important germ cell-specific role during ProSpg development. Mitotic stage (M) ProSpg are characterized as proliferative between E13.5 and E16.5, at which they then progress to transitional 1 (T1) ProSpg and enter a period of cellular quiescence. Soon after birth, T1 ProSpg become transitional 2 (T2) ProSpg by reentering the cell cycle until they become Spg at approximately postnatal day (P) 8 [22]. Recent studies by Law et al., 2019, suggest that a subpopulation of T1 ProSpg acquires high *Id4* expression, which begins proliferating to establish a robust initial pool of SSCs [23]. However, the transcriptional logic underlying these ProSpg transitions remains unknown.

Here we tested if and how TAF4b regulates critical transcriptional programs and cell states during early ProSpg development. First, we examined *Taf4b* expression in published single-cell (sc) RNA-seq data and confirmed its enriched expression in T1 ProSpg. Next, we performed bulk RNA-seq on *Taf4b*-deficient T1 ProSpg to reveal cell cycle and chromatin structure as top gene ontology (GO) categories modulated in the absence of TAF4b. We then determined that *Taf4b*-deficient ProSpg struggle to enter characteristic quiescence, noting significant perturbation around E14.5. For a genomic localization analysis of TAF4b in T1 ProSpg, we employed cleavage under targets and release using nuclease (CUT&RUN) and identified TAF4b peaks just upstream of transcription start sites (TSSs). Distinctive GO categories from the CUT&RUN included RNA processing, chromatin modification, and cell cycle regulation. The most consistent DNA motifs within the TAF4b-bound promoters included Sp/Klf family and NFY binding sites; a remarkable similarity to our findings in embryonic mouse oocytes, and for the first time linking these ubiquitous factors to the regulation of ProSpg transcription [24]. Finally, we compared our male embryonic germ cell data with our recently reported TAF4b genomic analyses of female embryonic germ cells revealing several core and sex-specific cellular processes regulated by TAF4b. Together these data suggest that TAF4b is a dynamic and vital integrator of the male germline transcription program during the transition of T1 ProSpg, required for proper mammalian spermatogenic development.

## RESULTS

### scRNA-seq reveals *Taf4b* mRNA expression peaks at P0 when proliferation markers are decreased

While we have observed that *Taf4b* mRNA expression peaks during mouse embryonic ProSpg development, single-cell sequencing data allow us to examine the gene expression within ProSpg at finer resolution, which we have reprocessed from published publicly available data sets [23,25,26]. In our analysis, we used the expression profile of *Taf4b* over time to evaluate what genes and biological processes are possibly modulated concurrently or downstream of TAF4b expression. After applying our computational workflow (see methods), the complete data set consisted of 17,310 germ cells in our uniform manifold approximation and projection (UMAP) (**Fig 1A)**. We then assessed the expression of *Taf4b* compared to its more ubiquitously expressed paralog, *Taf4a*. Across all time points, *Taf4a* mRNA expression was notably lower than *Taf4b* mRNA expression (**Fig 1B**). When observing the expression of *Taf4b* mRNA over the E12.5-P7 time course, we detected that *Taf4b* expression peaks at P0 after significantly low *Taf4b* expression at E12.5 (**Fig 1B**). The dynamic range of *Taf4b*-expressing cells across time suggests *Taf4b* may act more akin to a dimmer switch than on/off regulation. Because previous experiments demonstrated that the *Taf4b*-deficient gonad exhibited changes in germ cell proliferation, we compared this newly characterized *Taf4b* expression profile to known markers of proliferation such as *Mki67*. We saw an inverse expression pattern of *Mki67* and *Taf4b*, where the significantly reduced expression of *Mki67* between E16.5-P0 points to the characteristic quiescent period of T1 ProSpg (**Fig 1B**). *Plk1* mRNA, another marker of proliferation, is decreased similarly from E16.5 to P2 (**Fig 1B**). As a positive control for the quality of this data set, the highest levels of the SSC marker gene *Id4* were detected from P0-P3 as previously shown in the publication of origin (**Fig 1B**; [23]). The combined decreased markers of proliferation and increased *Taf4b* expression led us to hypothesize that *Taf4b* may play a role in the cell cycle transitions during ProSpg development.

**Figure 1.**
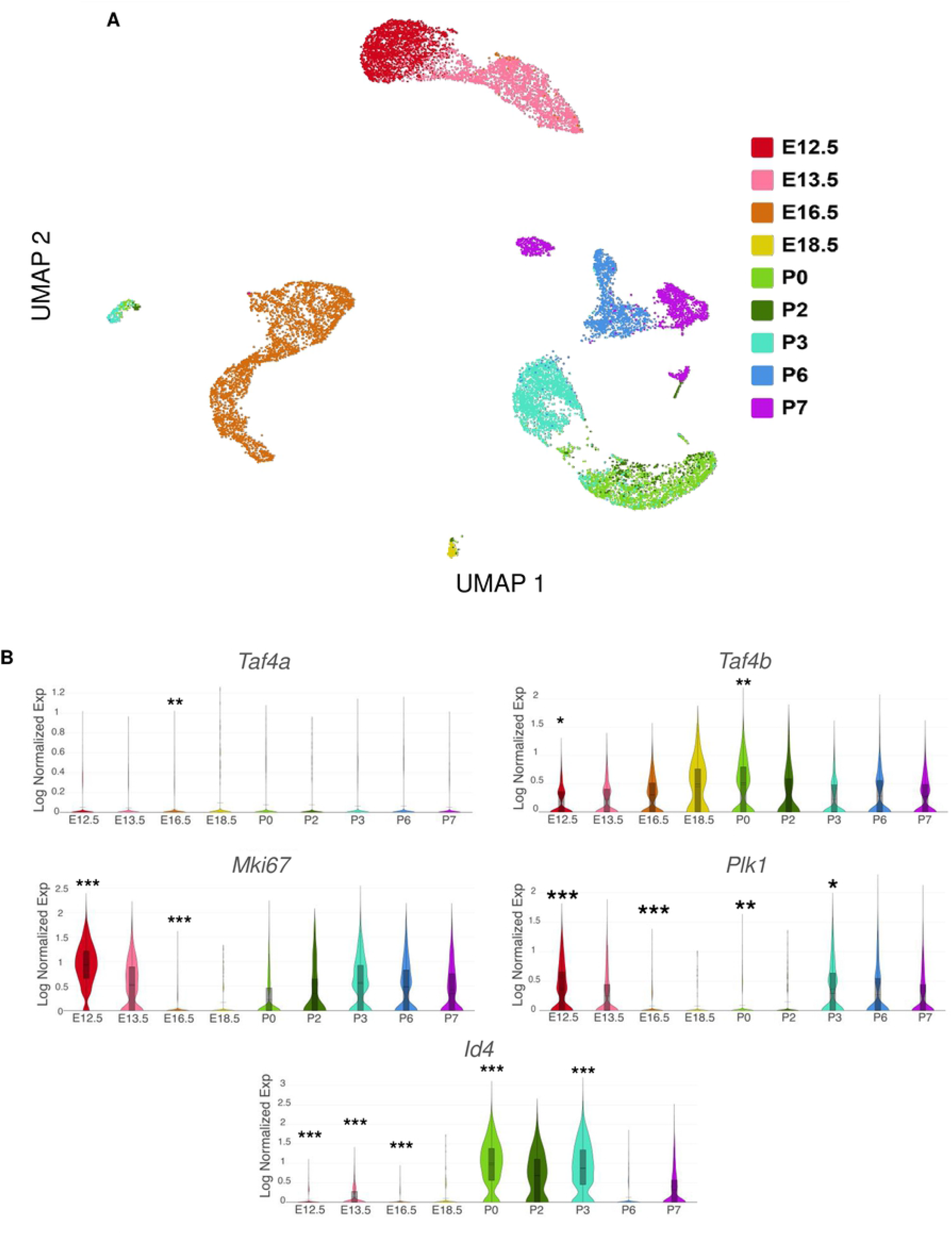
Characterizing *Taf4b* mRNA expression across male germ cell development in mouse. (A) scRNA-seq UMAP of E12.5-P7 germ cells colored by time point. (B) Expression of *Taf4a, Taf4b, Mki67, Plk1*, and *Id4* across the developmental time course. P-values are denoted (*** = p < 0.001, ** = p < 0.01, * = p < 0.05) and are adjusted using the Benjamini-Hochberg correction method for multiple tests.

### Global RNA-seq reveals *Taf4b*-affected gene expression programs in E16.5 ProSpg

Given our previous finding that proper postnatal germ cell expansion was disrupted in the absence of *Taf4b* [18], we were surprised to observe an increase in *Taf4b* mRNA during the expected ProSpg transition to quiescence. We thus aimed to determine the transcriptional changes resulting from *Taf4b’s* absence in E16.5 male germ cells. In order to understand the transcriptomic differences between *Taf4b*-wildtype, -heterozygous, and -deficient Pro-Spg, we used mice that were transgenic for *Oct4-eGFP* to sort for GFP^+^ germ cells and performed RNA-seq at E14.5 and E16.5. At E14.5, M ProSpg are expected to still be proliferative but by E16.5 most have transitioned to T1 ProSpg and have entered a period of quiescence. At E14.5, we performed RNA-seq on 8 samples (4 *Taf4b*-wildtype and 4 *Taf4b*-deficient) from 4 different collection dates (**Table S1**). The resulting principal component analysis (PCA) plot shows that the samples largely group together based on collection date and suggesting that collection date 1 was an outlier (**Fig S1A**). To compensate for this, we compared the *Taf4b*-wildtype and *Taf4b*-deficient samples which had a littermate of the other genotype (therefore excluding samples from collections 2 and 4) and corrected for the influence of litter collection dates using DESeq2. We identified 178 differentially expressed genes (DEGs) between *Taf4b*-wildtype and -deficient ProSpg, which were defined as protein-coding, average transcripts per million (TPM) expression > 1, and adjusted p-value < 0.05 (**Fig S1B, Table S1**). From this list of DEGs, 170 were increased in *Taf4b*-deficient ProSpg and 8 were decreased in *Taf4b*-deficient ProSpg. When we performed GO analysis of all the E14.5 DEGs, multiple development categories were enriched, including “gonad development” and “muscle tissue development” (**Fig S1C**) and increased DEGs from Figure 2B fall into one or more of these categories. While interesting, we found that our E16.5 RNA-seq data set on *Taf4b*-wildtype, -heterozygous, and -deficient ProSpg was more informative (**Table S2**). The PCA plot again indicated that there was grouping of samples due in part to the collection date (**Fig 2A**). Because *Taf4b*-heterozygous mice are fertile and there was only one DEG between *Taf4b*-wildtype and -heterozygous samples, we grouped the wildtype and heterozygous samples together and compared this group to the *Taf4b*-deficient ProSpg. We found 2751 DEGs between the *Taf4b*-wildtype/-heterozygous and -deficient ProSpg, which were defined as protein-coding, average TPM expression > 1, and adjusted p-value < 0.05 (**Fig 2B**). From this list of DEGs 1510 were increased in *Taf4b*-deficient ProSpg and 1241 were decreased in *Taf4b*-deficient ProSpg. Some notable genes that were among the DEGs were *Id4, Tex19*.*2*, and *Mki67*.

**Figure 2.**
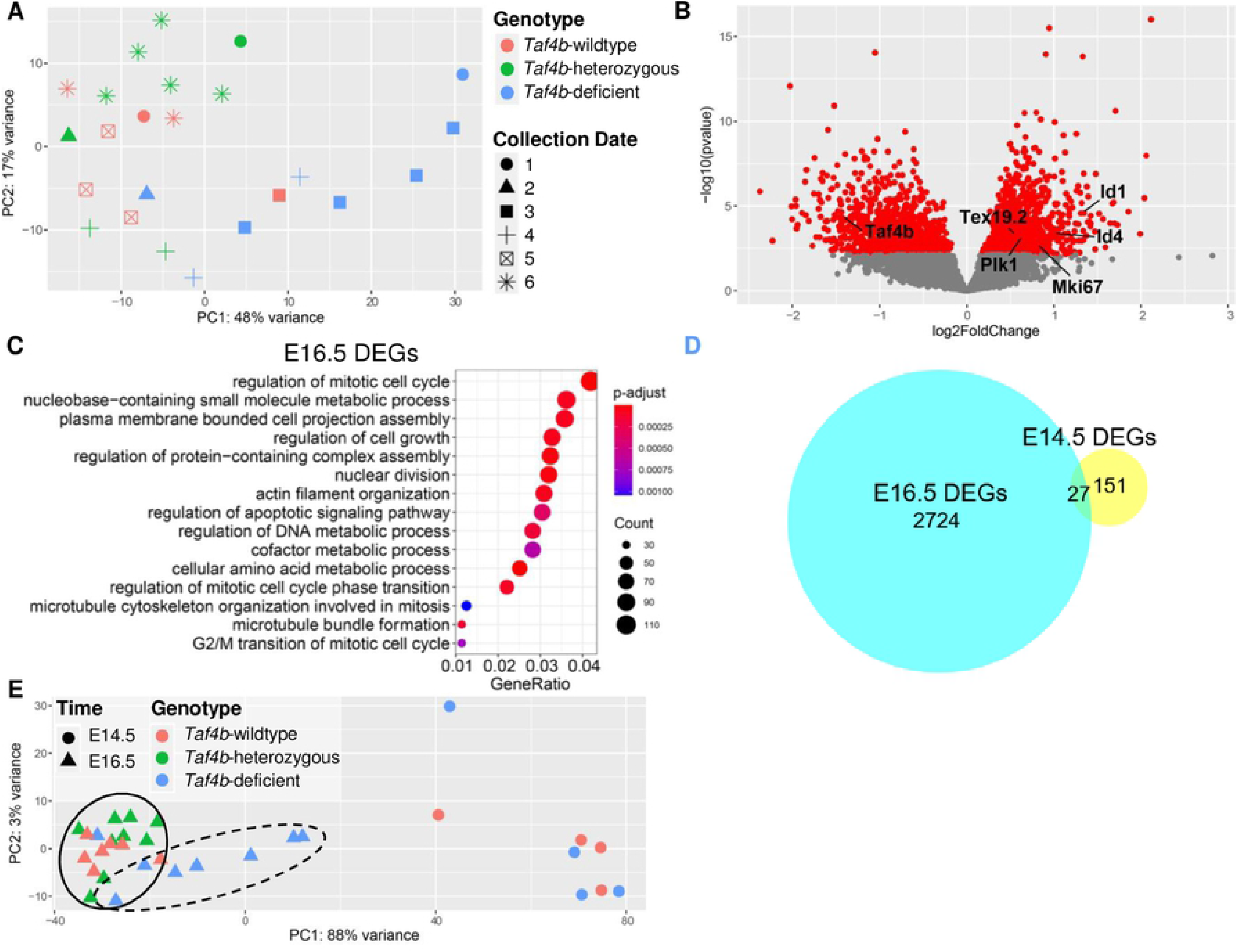
E16.5 ProSpg RNA-seq experiment reveals significant regulation of mitotic gene programs. (A) PCA plot of the E16.5 samples labeled based on *Taf4b* genotype and collection date. (B) Volcano plot of genes, the significant genes (protein-coding, p-adj < 0.05, avg TPM > 1) are labeled in red and DEGs of interest plus *Taf4b* are specified. (C) Dotplot of GO biological process analysis of DEGs from (B). (D) Venn diagram of DEGs from E14.5 and E16.5 RNA-seq experiments. (E) PCA plot of E14.5 and E16.5 samples labeled based on *Taf4b* genotype and age.

To assess *Taf4b*-dependent pathways, we performed GO analysis of all the E16.5 DEGs and found numerous mitotic categories enriched, including “regulation of mitotic cell cycle” and “G2/M transition of mitotic cell cycle” (**Fig 2C**). Examining the protein-protein interactions between the top 1500 DEGs revealed major nodes at *Trp53*, a tumor suppressor gene, and another node containing several cell cycle genes including *Plk1* and *Cdk1* (**Fig S2**). We then compared our E16.5 RNA-seq data to the E14.5 data and found that only 27 genes were shared between the two datasets and consisted of the genes: *Actg2, Adamts1, Armcx3, Atp1b2, Capn6, Cd200, Col14a1, Col16a1, Col5a2, Col6a1, Ctsh, Epdr1, Grb10, H2bc4, Magea8, Mef2c, Nrxn3, Prprs2, Prrx1, Sema4a, Sgce, Sorcs2, Sox11, Spp1, Taf4b, Tex19*.*2*, and *Tie1* (**Fig 2D**). We then examined our E14.5 and E16.5 RNA-seq datasets on the same PCA plot and noticed that the E16.5 *Taf4b*-deficient samples (dotted line) were closer to the E14.5 samples on the PC1 axis than their wildtype and heterozygous counterparts (solid line) (**Fig 2E**). This interestingly suggests that the E16.5 *Taf4b*-deficient ProSpg may have more similarities to E14.5 ProSpg than wildtype or heterozygous have to E14.5 ProSpg. Due to significant X chromosome expression effects we observed in *Taf4b*-deficient E16.5 female germ cells [24], we examined key aspects of X chromosome dosage to see if similar effects were occurring in *Taf4b*-deficient male germ cells. We investigated if there was a significant difference in autosomal expression compared to X chromosome expression, the X:A ratio, and the relative X expression (RXE) in E14.5 and E16.5 male germ cells (**Fig S3**). No significant difference in any calculation was observed, suggesting that these significant X chromosome effects are unique to *Taf4b*-deficient female germ cells [24].

### *Taf4b*-deficient ProSpg are delayed in their entry into to quiescence

Mouse M phase ProSpg enter a characteristic period of quiescence between E13.5 and E16.5. Based upon our scRNA-seq and bulk RNA-seq data, we hypothesized that TAF4b could promote the entry and/or maintenance of ProSpg quiescence. We performed indirect immunofluorescence on *Taf4b*-wildtype and *Taf4b*-deficient testis sections to detect TRA98 (a germ cell marker) and KI67 from E13.5 to E18.5. Representative images of each genotype and time point are shown (**Fig 3A**), as well as the quantification of the KI67-negative (KI67^-^) quiescent wildtype and *Taf4b*-deficient ProSpg (indicated by TRA98 signal) at each timepoint (**Fig 3B**). At E13.5, the majority of M ProSpg were still proliferating as indicated by the nuclear localization of KI67; occurrences are labeled with the yellow arrows. (**Fig 3A**). At E14.5, there is a significant reduction of ProSpg entering quiescence in the *Taf4b*-deficient compared to wildtype controls (**Fig 3B**). Interestingly, the number of quiescent cells at both E16.5 to E18.5 did not differ between controls and *Taf4b-*deficient Pro-Spg. Together, these data indicate that although *Taf4b*-defcient ProSpg can largely achieve quiescence, they struggle with its timely entry and transition.

**Figure 3.**
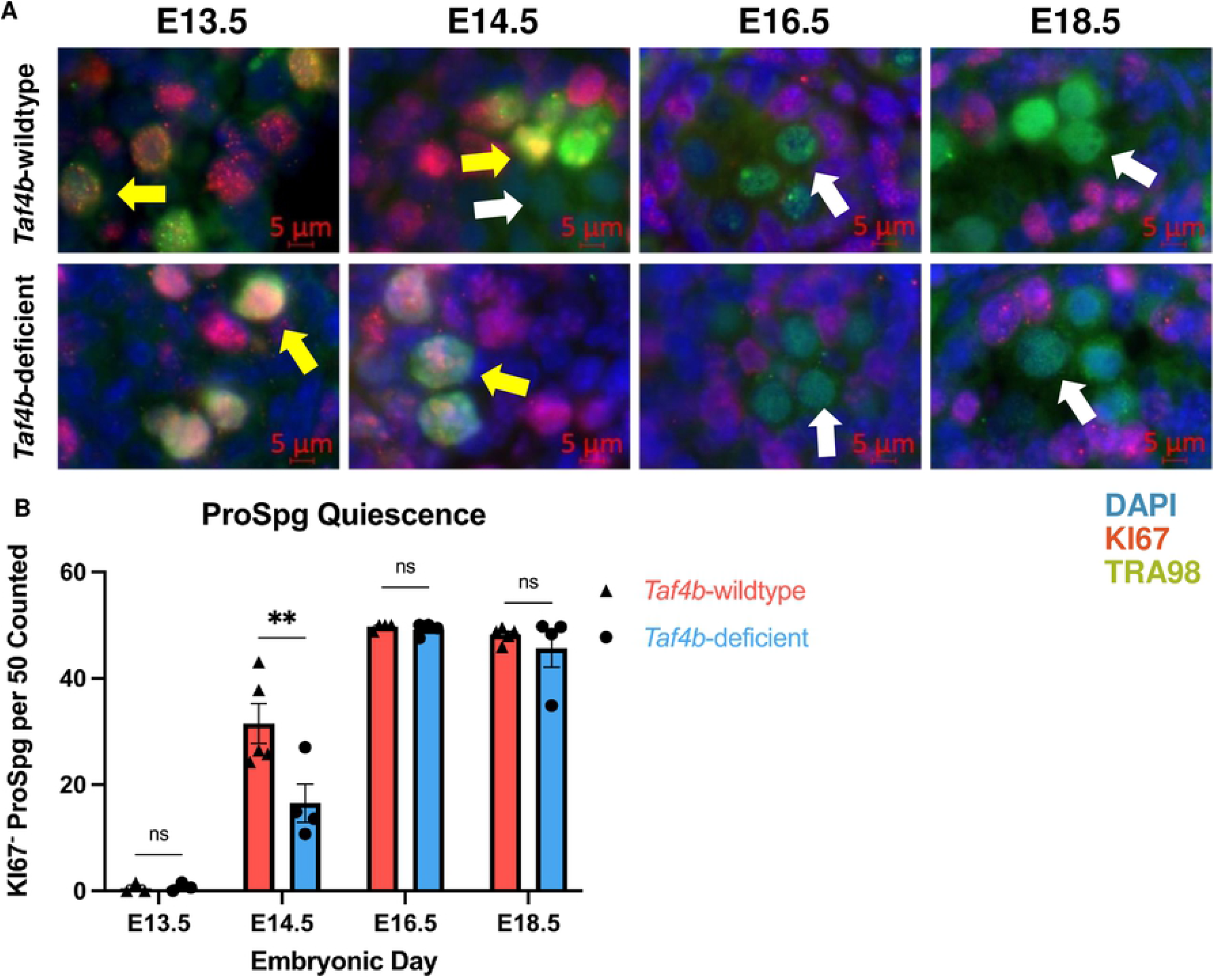
*Taf4b*-deficient ProSpg have increased KI67 at E14.5. (A) Immunofluorescence of E13.5-E18.5 testis sections comparing *Taf4b*-wildtype to *Taf4b*-deficient ProSpg. Antibodies for TRA98 (green), KI67 (red) were used and DAPI labels DNA. Colocalization of KI67 and TRA98 can be seen as yellow fluorescence and is labeled with yellow arrows, while the absence of KI67 is marked by the white arrows. (B) Quantification of KI67^-^/TRA98^+^ cells in *Taf4b*-wildtype (green) and *Taf4b*-deficient (red) testis sections. *Taf4b*-deficient ProSpg have significantly fewer KI67^-^ ProSpg than wildtype at E14.5 (n=3-5 mice, ** = adjusted p-value < 0.01 as determined by two-way ANOVA, ns = not significant).

### CUT&RUN identifies putative direct targets of TAF4b in E16.5 ProSpg

To distinguish which DEGs identified in our E16.5 bulk RNA-seq experiment were direct targets of TAF4b, we performed Cleavage Under Targets and Release Using Nuclease (CUT&RUN), a genome mapping technique to identify enrichment of specific proteins and histone modifications in the genome [27,28]. We isolated two replicates of wildtype *Oct4*-eGFP E16.5 male germ cells using FACS and examined the genomic localization of TAF4b. H3K4me3 served as a positive control and marked active promoters, and IgG served as a negative control. CUT&RUN data analysis using the program Homer identified 64,891 H3K4me3 peaks and 7,861 TAF4b peaks in Replicate 1 and 1,730 H3K4me3 peaks and 848 TAF4b peaks in Replicate 2 (**Table S3**). We also found that 73% and 88% of TAF4b peaks were classified as localizing to promoters/transcription start site (“Promoter-TSS”) for Replicate 1 and 2, respectively (**Fig 4A**). Of all the TAF4b peaks, 617 overlapped between the replicates (**Fig 4B**). When plotting the enrichment profile of TAF4b relative to TSSs, we found the highest TAF4b enrichment upstream of the TSS (**Fig 4C**). To more closely examine the localization of TAF4b signal near TSSs, we plotted the distance of TAF4b “promoter-TSS” peaks from both replicates to the TSS (**Fig 4D**). There is strong enrichment of TAF4b peaks between -200 bp to + 50 bp from the TSS, with the highest number of TAF4b peaks located at -20 to 0 bp away from the TSS. When we performed GO analysis of the shared TAF4b-bound gene promoter-TSSs between the two replicates, we found categories related to mRNA processing, RNA splicing, cell cycle, Wnt signaling, and DNA recombination/replication (**Fig 4E**).

**Figure 4.**
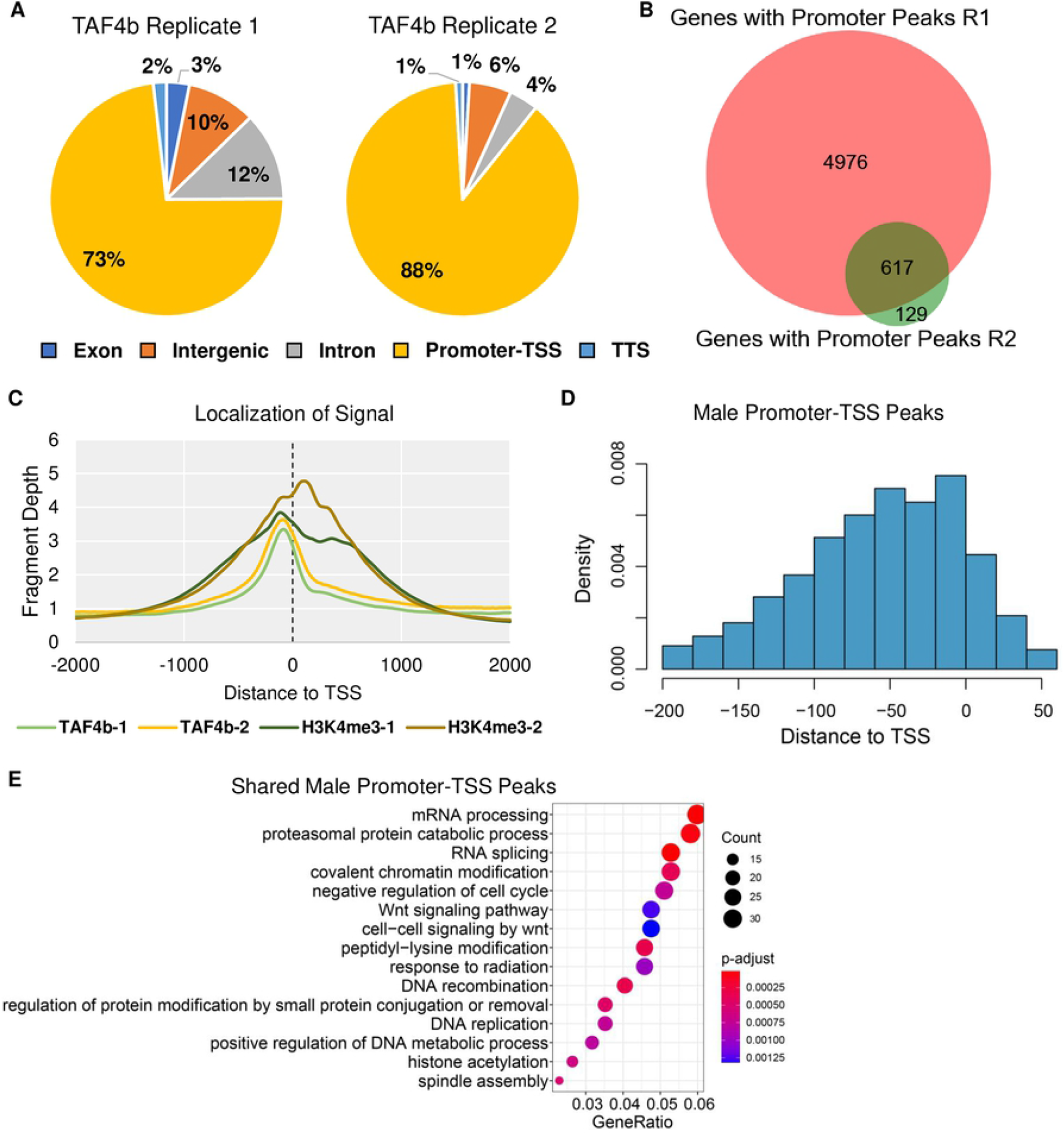
E16.5 ProSpg CUT&RUN identifies direct targets of TAF4b. (A) Pie charts of TAF4b peak locations in male germ cell CUT&RUN replicates. (B) Venn diagram of “promoter-TSS” peaks shared between the CUT&RUN replicates. (C) Average enrichment of TAF4b and H3K4me3 signal near TSSs (dotted line) for each replicate. (D) Histogram of TAF4b “promoter-TSS” peaks in relation to the TSS. (E) GO biological process dotplot for the shared CUT&RUN peaks categorized as “promoter-TSS”.

This CUT&RUN experiment also allowed us to begin inquiring as to which DEGs identified in our RNA-seq experiment were putative direct targets of TAF4b. When comparing our DEGs to “promoter-TSS” peaks of TAF4b we found 876 DEGs that had at least one peak near their TSS (**Fig 5A**). GO analysis of these peaks found that the biological process category most notably enriched in these data was “regulation of the mitotic cell cycle” (**Fig 5B**). A volcano plot of these TAF4b-bound E16.5 DEGs revealed far more upregulated DEGs (genes higher in *Taf4b*-deficient germ cells relative to wildtype and heterozygous, 674 genes) than downregulated DEGs in *Taf4b*-deficient ProSpg (202 genes) (**Fig 5C**). As examples of TAF4b-bound DEGs, we present tracks for *Nfya, Prc1, H2afx, Klf6*, and *Cdk20* (**Fig 5D-E**). *Nfya* is a component of the NFY protein complex and interestingly TAF4b peaks were located between the TSS of *Nfya* and another gene *Oard1* which was not a DEG. *Prc1* encodes a microtubule-binding protein that plays a role in mitosis and binds to another gene important to mitosis, *Plk1* [29]. *H2afx* encodes an essential DNA damage response protein and has been found in our previous research to be increased in *Taf4b*-deficient oocytes [30]. *Klf6* encodes a transcription factor that plays a role in several developmental processes such as cell proliferation and differentiation [31]. *Cdk20*, decreased in *Taf4b-*deficient ProSpg, encodes a positive regulator of the cell cycle by activating CDK2 and cyclin D [32]. Many other TAF4b-bound genes of interest were identified in only one replicate or were not a DEG in the E16.5 RNA-seq data (**Fig S4**). A protein-protein interaction network of these 876 TAF4b-bound DEGs revealed *Cdk1* and other mitosis-related genes as a major group of associated genes and interestingly the entire NFY complex (*Nfya, Nfyb*, and *Nfyc*) was present (**Fig S5**).

**Figure 5.**
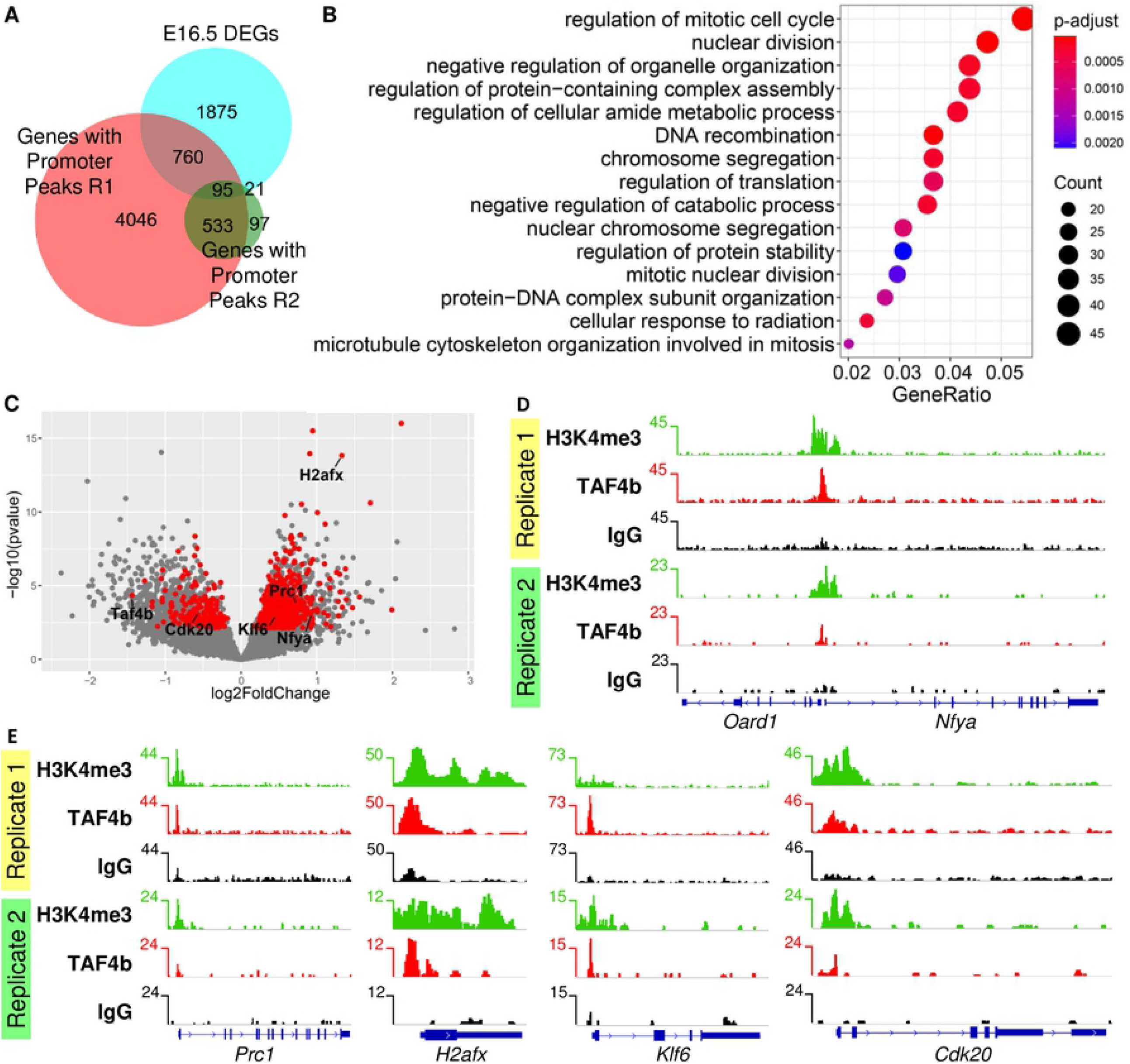
E16.5 CUT&RUN identifies putative direct targets of TAF4b in germ cells. (A) Venn diagram of CUT&RUN promoter peaks and RNA-seq DEGs. (B) Biological process GO dotplot of the 876 genes that are in the list of DEGs and had a TAF4b promoter-TSS peak in at least one of the two germ cell samples. (C) Volcano plot of the 876 DEGs that had at least one TAF4b promoter peak (red dots). (D) Gene track of *Oard1* and *Nfya*, in which *Nfya* was a DEG that had a TAF4b promoter-TSS called in both replicates. (E) Gene tracks of *Prc1, H2afx, Klf6*, and *Cdk20*, which were DEGs that had a TAF4b promoter-TSS called in both replicates.

We then explored the motifs associated with TAF4b-enriched peaks in our CUT&RUN data and found consistent enrichment of CCAAT and GC motifs associated with the NFY complex and the Sp/Klf transcription factor family, respectively. This enrichment was shared between the two CUT&RUN replicates, the DEGs that contained at least one TAF4b peak (both upregulated and downregulated DEGs), and from all the DEGs that contained at least one TAF4b peak (**Fig S6A-D&F**). When performing motif enrichment on “promoter-TSS” peaks based on distance to TSS, we found that Sp1 was almost always the top motif and there was interesting enrichment of the Ets family motifs from 0 bp to +50 bp away from the TSS (**Fig S6E**). As previously noted the highest number of TAF4b “promoter-TSS” peaks were located -20 to 0 bp away from the TSS (**Fig 4D**), which is approximately around where we might expect TAF4b enrichment. To further explore these unexpected motifs, we created boxplots (no outliers included) to display the distance of TAF4b “promoter-TSS” peaks from both replicates, all TAF4b “promoter-TSS” peaks for genes that were also DEGs, TAF4b “promoter-TSS” peaks for genes that were only Downregulated DEGs, and TAF4b “promoter-TSS” peaks for genes that were only Upregulated DEGs (**Fig S6G**). Their median locations from the TSS were -53 bp, -51 bp, -48.5 bp, and -53 bp, respectively, with no significant differences between the groups. These data combining RNA-seq and CUT&RUN of TAF4b suggest that TAF4b directly regulates RNA processing and chromatin genes through an unexpected function of TAF4b through Sp/Klf family motifs. However, more canonical functions of TAF4b cannot be ruled out, as “TATA-Box (TBP)/Promoter” did appear as a significantly enriched motif in both replicates, it was ranked 109 in Replicate 1 and 147 in Replicate 2.

### Integrating E16.5 ProSpg and oocyte genomic data reveals sex-independent and sex-dependent pathways of TAF4b regulation

Our work indicates that TAF4b performs essential developmental functions in spermatogenesis and oogenesis. We thus compared our findings in E16.5 male ProSpg (from this study) to E16.5 oocytes [24]. We plotted all our samples in one PCA plot and observed that the samples separated strongly by age and sex (**Fig 6A**). When comparing DEGs, there were 177 DEGs shared between E16.5 female DEGs and E14.5 and E16.5 male DEGs (**Fig 6B**). The only significant biological process GO category for these 177 DEGs was “cellular process involved in reproduction in multicellular organism”, which included these 14 genes: *Nobox, Ercc1, Spag6l, Dpep3, Piwil4, Slc26a6, Alms1, Psme4, Aurka, Mybl1, Plk1, Chtf18, Lin28a*, and *Tnk2* (**Fig 6C**). For the DEGs unique to the E16.5 ProSpg, we found categories including “regulation of mitotic cell cycle” and “actin filament organization” (Fig 6D). For the DEGs unique to the E16.5 oocytes, we found categories including “covalent chromatin modification” and “microtubule-based movement” (**Fig 6E**). When combining our male CUT&RUN replicates with the female CUT&RUN TAF4b peaks, we found 196 “promoter-TSS” peaks found in all four replicates and 611 genes that had three peaks (**Fig 6F**). The 196 genes were primarily associated with mRNA processing and splicing (**Fig 6G**) while the 611 genes also included mRNA processing plus DNA repair and cell cycle categories (**Fig 6H**). These might be the core set of genes that TAF4b directly regulates. Accordingly, we conclude from all of these data that TAF4b regulates both sex-independent and sex-dependent embryonic germ cell development further highlighting its critical transcription functions regulating male and female mammalian fertility.

**Figure 6.**
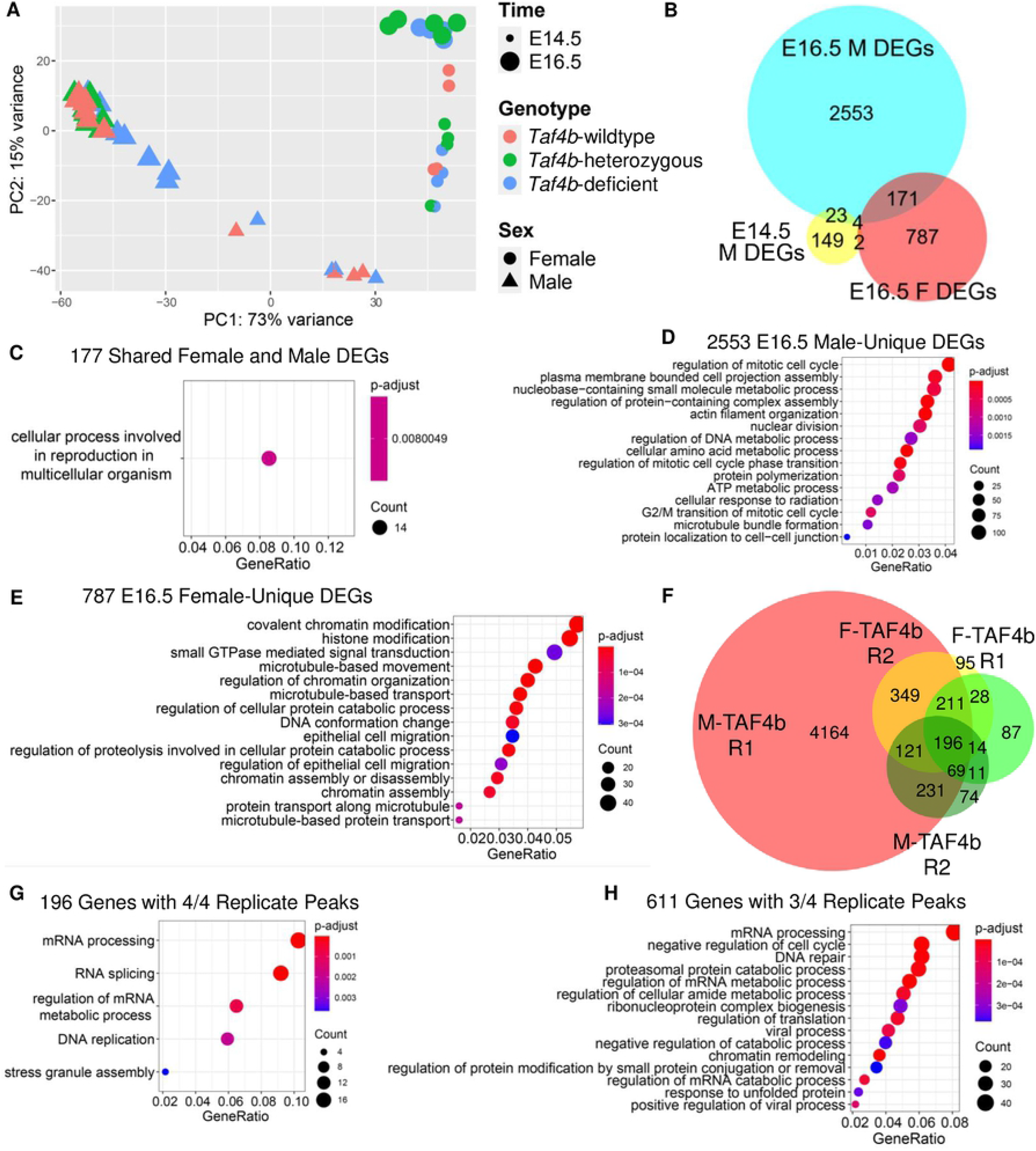
Male versus female data. (A) PCA plot of all E14.5 and E16.5 RNA-seq samples. (B) Venn diagram of DEGs from three separate RNA-seq experiments in embryonic germ cells. (C) Dotplot of GO biological process for the 171 genes shared between E16.5 male DEGs and E16.5 female DEGs. (D) Dotplot of GO biological process categories for the 2553 DEGs that were unique to E16.5 *Taf4b*-deficient prospermatogonia. (E) Dotplot of GO biological process categories for the 787 DEGs that were unique to E16.5 *Taf4b*-deficient oocytes. (F) Venn diagram of all genes with promoter peaks from all E16.5 female and male CUT&RUN replicates. (G) Dotplot of GO biological process categories for the 196 genes that have a promoter peak in all four CUT&RUN replicates. (H) Dotplot of GO biological process categories for the 611 genes that have a promoter peak in at least three of the replicates.

## DISCUSSION

Soon after sex specification in mammals, embryonic female and male germ cells receive distinct signals to initiate a sex-specific germ cell differentiation program. However, it is unclear how these newly defined germ cells integrate signals to execute precise and sex-specific gene expression programs and initiate the process of producing eggs and sperm as mature adults. We have discovered that the embryonic germline-enriched TFIID subunit, TAF4b, is required for male and female germ cell development and maintenance in mice. Here we identify many ProSpg genes whose expression are affected by the loss of TAF4b and whose promoter regions display TAF4b occupancy just upstream of the TSS. TAF4b enrichment at ProSpg core promoters correlates with the binding sites for Sp1 and NFY, two well-characterized and ubiquitous promoter-proximal and sequence-specific transcription factors. The integration of these high throughput genomic sequencing and mapping data indicate that TAF4b promotes male-specific cell cycle gene expression required for the timely entry of ProSpg into quiescence. Interestingly, TAF4b regulates genes shared between male and female embryonic germ cells at identical time points despite the two dramatically different germ cell trajectories, i.e., mitotic quiescence vs. meiosis. The similitude further indicates that it plays critical roles in both sex-dependent and independent embryonic germ cell identity at equivalent chronological time points.

Several notable regulators of post-transcriptional gene expression play similar developmental functions as TAF4b when ProSpg transition towards quiescence. The most notable is the RNA-binding protein DND1 which displays intriguing parallels with the functions of TAF4b shown here. DND1 deficiency in mice leads to defects of prospermatogonial entry into quiescence and increased germ cell loss [33]. However, the Ter mutation in the *Dnd1* gene on the 129 genetic background strikingly results in spontaneous teratoma formation at E16.5 [33]. While *Taf4b*-deficient testes display similar kinetics in male germ cell loss and fertility disruption, our *Taf4b*-deficiency presented here is on a C57Bl/6 background where we might not expect to see these teratomas [17]. Strikingly, TAF4b and DND1 share common regulation of mitotic cell cycle and chromatin modification gene expression, suggesting they are two distinct modes of regulating a common prospermatogonial quiescence and survival program ([34], this study). A second RNA-binding protein, NANOS2, is also responsible for prospermatogonial development and has been recently shown to be bound to and work with DND1 in regulating the specific RNA loading into the CNOT deadenylase complex [34,35]. The developmental similarities of perturbed quiescence, increased germ cell loss, and infertility in *Nanos2, Dnd1*, and *Taf4b* mouse mutants suggest that regulating the gene expression of prospermatogonia transitions is critical to properly set up the future cells that arise from this lineage in the postnatal testis.

There are also several key transcription factors that direct cell cycle progression and transcription during prospermatogonial development. The master regulator of the G1 phase of the cell cycle, RB1, is required for the timely mitotic arrest of early prospermatogonia [36]. A recent germ cell-specific knockout approach for RB1 linked this cell cycle defect to disrupting the metabolic nature of these developing prospermatogonia and their corresponding ability to become SSCs [37]. While the timing of RB1 and TAF4b functions in promoting quiescence are similar, RB1 deficiency leads to much more severe cell cycle defects than *Taf4b*-deficiency, suggesting they play more complementary roles in SSC development [37]. RHOX10 is a homeodomain sequence-specific DNA-binding protein found in the reproductive homeobox cluster of the X chromosome and is required for prospermatogonial and SSC development. A recent genomic study using similar tools implemented here uncovered a network of transcription factors that are direct targets of RHOX10, most notably DMRT1 and ZBTB16 [38]. While there are minor differences in the timing and cells used in these two studies, Tan et al. uncovered a critical CCAAT box bound by RHOX10 in the Dmrt1 promoter and predicted it to be a binding site for the NFY transcription factor, which was the most correlated binding site at TAF4b bound core promoters ([8], this study). It will be interesting to determine how these transcriptional regulators work together and separately to promote prospermatogonial development required for the proper establishment of SSCs and long-term mammalian spermatogenesis.

While we focus here on TAF4b’s role in promoting the early development of the male germline, we have recently reported that at similar embryonic time points in development, TAF4b plays a critical role in early oocyte differentiation [24]. As female and male embryonic germ cells express unique sets of genes and navigate different biological processes, i.e., meiotic initiation vs. mitotic arrest, it is surprising that the same transcription factor would have distinct functions in the early life of these future gametes. In fact, we see both shared and distinct promoters bound by TAF4b when we compare fine-scale genomic mapping for TAF4b between female and male E16.5 embryonic germs cells by CUT&RUN. We propose that one joint function of TAF4b, and its associated regulatory complex, is to promote the embryonic germ cell identity and survival after PGC specification, analogous to the germ cell licensing function of DAZL [39]. In addition, TAF4b has likely evolved sex-specific functions that allow it to promote context-specific transcription events in both male and female developing germ cells. One commonality of these two populations is that even though they enter meiosis at different times, they both exit the mitotic cell cycle at the end of PGC development, and TAF4b, along with many other regulatory proteins, may help direct this transition. Finally, genome-wide epigenetic modifications are erased in both female and male germ cell populations at the end of PGC development. In the context of a more open genome, TAF4b and related regulators could help maintain the appropriate chromatin environment for controlling transcription-coupled DNA repair mechanisms unique and critical to the germline genome of both sexes and, ultimately, the next generation.

## MATERIALS AND METHODS

### Ethics statement

This study was approved by Brown University IACUC protocol #21-02-0005. The primary method of euthanasia is CO_2_ inhalation and the secondary method used is cervical dislocation both as per American Veterinary Medical Association (AVMA) guidelines on euthanasia.

### Mice

Mice, homozygous for an *Oct4-eGFP* transgene (The Jackson Laboratory: B6;129S4-*Pou5f1*^*tm2Jae*^/J), were mated for CUT&RUN collections. Mice, homozygous for an *Oct4-eGFP* transgene (The Jackson Laboratory: B6;129S4-*Pou5f1*^*tm2Jae*^/J) and heterozygous for the *Taf4b*-deficiency mutation (in exon 12 of the 15 total exons of the *Taf4b* gene that disrupts the endogenous *Taf4b* gene), were mated for mRNA collections. Timed matings were estimated to begin at day 0.5 by evidence of a copulatory plug. The sex of the embryos was identified by confirming the presence or absence of testicular cords. Genomic DNA, isolated from tail biopsies using Qiagen DNeasy Blood & Tissue Kits (Cat #: 69506), was used for PCR genotyping assays. All animal protocols were reviewed and approved by Brown University Institutional Animal Care and Use Committee and were performed in accordance with the National Institutes of Health Guide for the Care and Use of Laboratory Animals. Gonads were dissected out of embryos into cold PBS.

### Immunofluorescence

Testes were gathered from embryonic mice (E13.5-E18.5) and paraffin-embedded. The Molecular Pathology Core at Brown University completed tissue preparation and performed sectioning. After the serial sections were prepared, the samples were permeabilized in 0.5% Triton X-100 for 25 minutes. They were then incubated first in blocking buffer (PBS with 3% goat serum, 1% BSA, and 0.5% Tween-20) and secondly in primary antibody prepared in blocking solution for 24 hours at 4°C. The samples were then washed in PBS+0.5% Tween-20 (PBST), incubated with secondary antibody for 1 hour at 37°C, washed in PBST, and mounted in DAPI-containing Vectashield Mounting Medium (Vector Laboratories H-1200, Burlingame, CA). Primary antibodies used were rat anti-TRA98 (1:100, abcam ab82527) and anti-KI67 rabbit mAb (1:100, Cell Signaling Technology D3B5 9129S). For immunofluorescence, the secondary antibodies used were Alexa Fluor 555-conjugated anti-rat IgG (1:500; abcam ab150158) and Alexa Fluor 488-conjugated anti-rabbit IgG (1:500; abcam ab150081). Images were obtained with a Zeiss Axio Imager.M1.

### Embryonic gonad dissociation and fluorescence-activated cell sorting

To dissociate gonadal tissue into a single-cell suspension, embryonic gonads were harvested and placed in 0.25% Trypsin/EDTA and incubated at 37°C for 15 and 25 minutes for E14.5 and E16.5 gonads, respectively, as previously described [21]. Eppendorf tubes were flicked to dissociate tissue halfway through and again at the end of the incubation. Trypsin was neutralized with FBS. Cells were pelleted at 1,500 RPM for 5 minutes, the supernatant was removed, and cells were resuspended in 100 μL PBS. The cell suspension was strained through a 35 μm mesh cap into a FACS tube (Gibco # 352235). Propidium iodide (1:500) was added to the cell suspension as a live/dead distinguishing stain. Fluorescence-activated cell sorting (FACS) was performed using a Becton Dickinson FACSAria III in the Flow Cytometry and Cell Sorting Core Facility at Brown University. A negative control of a non-transgenic mouse gonad was used for each experiment to establish an appropriate GFP signal baseline. Dead cells were discarded, and the remaining cells were sorted into GFP^+^ and GFP^-^ samples in PBS at 4°C for each embryo.

For RNA-seq analysis, GFP^+^ cells from each individual embryo were kept in separate tubes and were then spun down at 1,500 RPM for 5 minutes, had PBS removed, and were then resuspended in Trizol (ThermoFisher # 1556026). If samples had roughly less than 50 µL of PBS in the tube, Trizol was added immediately. The number of cells for each sample can be found in **Table S4**. Samples were stored at -80°C.

For CUT&RUN, germ cells from all the gonads were pooled prior to FACS. Sorted cells were then spun down at 1,500 RPM for 5 minutes and were resuspended in 300 µL of PBS, then split into three Eppendorf tubes. These three tubes of germ cells were then used forCUT&RUN. The number of cells for each sample were as follows: Replicate 1 germ cell samples had approximately 56,000 cells per tube (obtained from 22 embryos) and Replicate 2 germ cell samples had approximately 131,000 cells per tube (obtained from 28 embryos).

### Single cell RNA-seq data analysis

All computational scripts regarding single cell RNA-seq (scRNA-seq) used in this publication are available to the public: https://github.com/mg859337/Gura_et_al._TAF4b_male_transcription. SRP194420, SRP158811, and SRP178196 were downloaded from NCBI SRA onto Brown University’s high-performance computing cluster at the Center for Computation and Visualization. We used Law et al. and Nguyen et al., which enriched for germ cells via FACS sorting. We then combined the data with Tan et al., who collected the whole testis, thus establishing our comprehensive observation window from E12.5 to P7. The fastq files were aligned using Cell Ranger (v 6.0.0) count and then aggregated using Cell Ranger aggr. The cloupe file created from Cell Ranger aggr was used as input for Loupe Cell Browser (v 5.0). We selected the germ cells by first assessing the log normalized expression of germ cell marker *Dazl* within the complete 71,584 cell data set. Then, using unbiased clustering, we eliminated somatic cell clusters based on their mean *Dazl* expression being less than 0.85. The data set underwent further quality control and filtering based on fitting the metrics of 12.5-15.8 log2-normalized unique molecular identifiers, 10.6-12.8 log2-normalized features, and lastly, less than 24% mitochondrial transcript ratio.

### RNA-sequencing

Embryonic germ cells resuspended in Trizol were shipped to GENEWIZ (GENEWIZ Inc., NJ) on dry ice. Sample RNA extraction, sample QC, library preparation, sequencing, and initial bioinformatics were done at GENEWIZ. RNA was extracted following the Trizol Reagent User Guide (ThermoFisher Scientific). Glycogen was added (1 µL, 10 mg/mL) to the supernatant to increase RNA recovery. RNA was quantified using Qubit 2.0 Fluorometer (Life Technologies, Carlsbad, CA, USA) and RNA integrity was checked with TapeStation (Agilent Technologies, Palo Alto, CA, USA) to see if the concentration met the requirements.

SMART-Seq v4 Ultra Low Input Kit for Sequencing was used for full-length cDNA synthesis and amplification (Clontech, Mountain View, CA), and Illumina Nextera XT library was used for sequencing library preparation. The sequencing libraries were multiplexed and clustered on a lane of a flowcell. After clustering, the flowcell was loaded onto an Illumina HiSeq 4000 according to manufacturer’s instructions. The samples were sequenced using a 2×150 Paired End (PE) configuration. Image analysis and base calling were conducted by the HiSeq Control Software (HCS) on the HiSeq instrument. Raw sequence data (.bcl files) generated from Illumina HiSeq were converted into fastq files and de-multiplexed using bcl2fastq (v. 2.17). One mismatch was allowed for index sequence identification.

### RNA-seq data analysis

All computational scripts regarding RNA-seq used in this publication are available to the public: https://github.com/mg859337/Gura_et_al._TAF4b_male_transcription. All raw fastq files were initially processed on Brown University’s high-performance computing cluster. Reads were quality-trimmed and had adapters removed using Trim Galore! (v 0.5.0) with the parameters – nextera -q 10. Samples before and after trimming were analyzed using FastQC (v 0.11.5) for quality and then aligned to the Ensembl GRCm38 using HiSat2 (v 2.1.0) [40,41]. Resulting sam files were converted to bam files using Samtools (v 1.9) [42].

To obtain TPMs for each sample, StringTie (v 1.3.3b) was used with the optional parameters -A and -e. A gtf file for each sample was downloaded and, using RStudio (R v 4.0.2), TPMs of all samples were aggregated into one comma separated (csv) file using a custom R script. To create interactive Microsoft Excel files for exploring the TPMs of each dataset: the csv of aggregated TPMs was saved as an Excel spreadsheet, colored tabs were added to set up different comparisons, and a flexible Excel function was created to adjust to gene name inputs. To explore the Excel files, please find the appropriate tab (named “Quick_Calc”) and type in the gene name of interest into the highlighted yellow boxes. There is an Excel file for each dataset analyzed as a supplementary table.

To obtain count tables, featurecounts (Subread v 1.6.2) was used [43]. Metadata files for dataset were created manually in Excel and saved as a csv. These count tables were used to create PCA plots by variance-stabilizing transformation (vst) of the data in DESeq2 (v 1.22.2) and plotting by ggplot2 (v 3.1.0) [44,45]. DESeq2 was also used for differential gene expression analysis, where count tables and metadata files were used as input. We accounted for the litter batch effect in our mouse germ cells by setting it as a batch parameter in DESeq2. For the volcano plot, the output of DESeq2 was used and plotted using ggplot2. DEG lists were used for ClusterProfiler (v 3.16.1) input to create dotplots of significantly enriched gene ontology (GO) categories for all DEGs. The highest-confidence protein-protein interactions were identified using STRING, with unconnected proteins not shown in the images [46].

For X chromosome analysis, boxplots of log2 fold change between the autosomes and X chromosomes used the output of DESeq2 as input, based on other publications comparing autosomal and X chromosome expression [47]. The X:A ratio was calculated using pairwiseCI (v. 0.1.27), a bootstrapping R package, after filtering genes for an average TPM > 1 [48,49]. The RXE was calculated using a custom R script based after filtering genes for an average TPM > 1 and adding pseudocounts for log transformation (log2(x+1)), based on other RXE publications [50,51]. Venn diagrams were created using BioVenn (Hulsen et al., 2008). All plots produced in RStudio were saved as an EPS file type and then opened in Adobe Illustrator in order to export a high-quality JPEG image.

### CUT&RUN

The CUT&RUN performed in E16.5 germ cells followed the protocol in Hainer and Fazzio, 2019. CUT&RUN antibodies were as follows: polyclonal rabbit TAF4b (as previously described [30]), monoclonal rabbit H3K4me3 (EMD Millipore # 05-745R), rabbit IgG (ThermoFisher# 02-6102), pA-MNase was a generous gift from Dr. Thomas Fazzio.

For library preparation, the KAPA HyperPrep kit (Roche Cat. No 07962363001) was used with New England Biolabs NEBNext Multiplex Oligos for Illumina (NEB #E7335). After library amplification through PCR, libraries were size selected through gel extraction (∼150-650 bp) and cleaned up using the Qiagen QIAquick Gel Extraction Kit (Cat. # 28704). CUT&RUN libraries in EB buffer were shipped to GENEWIZ (GENEWIZ Inc., NJ) on dry ice. Sample QC, sequencing, and initial bioinformatics were done at GENEWIZ.

The sequencing libraries were validated on the Agilent TapeStation (Agilent Technologies, Palo Alto, CA, USA), and quantified by using Qubit 2.0 Fluorometer (Invitrogen, Carlsbad, CA) as well as by quantitative PCR (KAPA Biosystems, Wilmington, MA, USA). The sequencing libraries were clustered on flowcells. After clustering, the flowcells were loaded on to the Illumina HiSeq instrument (4000 or equivalent) according to manufacturer’s instructions. The samples were sequenced using a 2×150bp Paired End (PE) configuration. Raw sequence data (.bcl files) generated from Illumina HiSeq were converted into fastq files and de-multiplexed using bcl2fastq (v. 2.20). One mismatch was allowed for index sequence identification.

### CUT&RUN data analysis

All computational scripts regarding CUT&RUN data analysis used in this publication are available at: https://github.com/mg859337/Gura_et_al._TAF4b_male_transcription and based on other CUT&RUN publications [28]. All raw fastq files were initially processed on Brown University’s high-performance computing cluster. Reads were quality-trimmed and had adapters removed using Trim Galore! (v 0.5.0) with the parameter -q 10 (https://www.bioinformatics.babraham.ac.uk/projects/trim_galore/). Samples before and after trimming were analyzed using FastQC (v 0.11.5) for quality and then aligned to the Ensembl GRCm39 using Bowtie2 (v 2.3.0). Fastq screen (v 0.13.0) was used to determine the percentage of reads uniquely mapped to the mouse genome in comparison to other species. Resulting sam files were converted to bam files, then unmapped, duplicated reads, and low quality mapped were removed using Samtools (v1.9). Resulting bam files were split into size classes using a Unix script. For calling peaks, annotating peaks, and identifying coverage around TSSs, Homer (v 4.10) was used [52]. For gene track visualization, the final bam file before splitting into size classes was used as input to Integrative Genomics Viewer (IGV) [53]. A custom genome was created using a genome fasta and gtf file for Ensembl GRCm39.

Pie charts were created using data from Homer output and Venn diagrams were created using BioVenn. Dotplots of Promoter-TSS peaks were made using ClusterProfiler. TSS plots were created using the “tss” function of Homer and plotted using Microsoft Excel. All plots produced in RStudio were saved as an EPS file type and then opened in Adobe Illustrator in order to export a high-quality JPEG image.

### Data availability

The male mouse E14.5 and E16.5 RNA-sequencing data are available from NCBI GEO under accession number GSE188351. The male mouse E16.5 CUT&RUN sequencing data are available from NCBI GEO under accession number GSE188701. The sequencing datasets accessed in this research are from the follow accession numbers: the scRNA-seq mouse data from E12.5 to P7 male sorted *Oct4-eGFP* gonads used for Figure 1 was obtained through NCBI GEO: GSE119045, GSE124904, and GSE130593.

## ACKNOWLEDGMENTS

We thank the Center for Computation and Visualization at Brown University for computational resources for scRNA-seq, RNA-seq, and CUT&RUN data analysis. We thank Kevin Carlson and the Brown University Flow Cytometry and Sorting Facility for expertise completing the flow sorting. The Brown University Flow Cytometry and Sorting Facility has received generous support in part by the National Institutes of Health (NCRR Grant No. 1S10RR021051) and the Division of Biology and Medicine, Brown University. As some of our insights were gained by reprocessing publicly available scRNA-seq datasets, we greatly appreciate both the researchers that generated and shared the data initially and the respective repositories for making them available. We are grateful to the NICHD/NIH for their generous support through awards 1F31HD097933, and 1R01HD091848 to MAG and RNF, respectively and thank the BSF for their generous support.

## Author Contributions

MAG, conception and design of RNA-seq and CUT&RUN experiments, collection and assembly of data, data analysis and interpretation, manuscript writing; MAB, conception and design of experiments, collection and assembly of data, data analysis and interpretation, manuscript writing; SR, collection and assembly of data; KMA, collection and assembly of data; KAS, mouse colony management, collection of gonadal tissue and cells, data analysis and interpretation, manuscript writing; RNF, conception and design of all experiments, data interpretation, manuscript writing and financial support.

## FIGURE LEGENDS

**Figure S1. E14.5 ProSpg RNA-seq experiment**. (A) PCA plot of the E14.5 samples labeled based on *Taf4b* genotype and collection date. (B) Volcano plot of genes with the significant genes (protein-coding, p-adj < 0.05, avg TPM > 1) labeled in red and DEGs of interest plus *Taf4b* are specified. (C) Dotplot of GO biological process analysis of DEGs.

**Figure S2. Protein-protein interactions of top 1500 E16.5 DEGs**. Protein-protein interactions graphic generated by STRING for the top 1500 E16.5 RNA-seq DEGs (highest confidence, unconnected nodes dropped) [46]. There was a significant enrichment of PPIs (p < 0.001).

**Figure S3. X chromosome expression at E14.5 and E16.5**. (A) E14.5 boxplots of log2 fold change values from DESeq2 of all genes comparing autosomal versus X chromosome log2 fold change (outliers removed), Welch’s t-test, NS = not significant. (B) E14.5 X:A ratio plot calculated through pairwiseCI after filtering for average TPM > 1 comparing Wt X:A ratio to *Taf4b*-deficient X:A Ratio. (C) E14.5 boxplots of relative X expression (RXE) calculations after filtering for average TPM > 1 and adding pseudocounts for log-transformation for each *Taf4b*-Wt and – deficient sample. (D) E16.5 boxplots of log2 fold change values from DESeq2 of all genes comparing autosomal versus X chromosome log2 fold change (outliers removed). (E) E16.5 X:A ratio plot calculated through pairwiseCI after filtering for average TPM > 1 comparing Wt and Het X:A ratio to *Taf4b*-deficient X:A Ratio. (F) E16.5 boxplots of relative X expression (RXE) calculations after filtering for average TPM > 1 and adding pseudocounts for log-transformation for each *Taf4b*-Wt/Het and –deficient sample.

**Figure S4. Selected gene tracks**. Genes tracks of *Sp1* (A) and *Foxo1* (B), which were TAF4b CUT&RUN “promoter-TSS” peaks shared between the replicates but not DEGs. Gene tracks of *Id4* (C) and *Tex19*.*2* (D), which were DEGs but had no TAF4b peaks called. (E) Gene track for *Plk1*, a DEG that had a TAF4b peak called in only Replicate 1. (F) Gene track for *Id1*, a DEG that had a TAF4b peak called in only Replicate 2. (G) Gene track for *Wnt2b*, a non-DEG that had no TAF4b peaks called but did have H3K4me3 peaks called in both replicates.

**Figure S5. Protein-protein interactions of TAF4b-bound DEGs identified by RNA-seq and CUT&RUN**. Protein-protein interactions graphic generated by STRING of DEGs from the E16.5 RNA-seq and had at least one TAF4b “promoter-TSS” peak (highest confidence, unconnected nodes dropped) [46]. There was a significant enrichment of PPIs (p < 0.0001).

**Figure S6. Strong motif consistency when examining subsets of E16.5 prospermatogonia CUT&RUN data**. (A) Top five TAF4b motifs from “promoter-TSS” peaks in male Replicate 1. (B) Top five TAF4b motifs from “promoter-TSS” peaks in male Replicate 2. (C) Top five TAF4b motifs from Upregulated DEGs that had at least one TAF4b “promoter-TSS” peak. (D) Top five TAF4b motifs from Downregulated DEGs that had at least one TAF4b “promoter-TSS” peak. (E) Diagram of the top three TAF4b “promoter-TSS” motifs in 50 bp windows relative to the TSS. (F) Top five motifs enriched at TAF4b “promoter-TSS” peaks for genes that were also DEGs, the promoter ID, and the associated p-value. (G) Boxplots (no outliers included) of peaks relative to the TSS for all TAF4b “promoter-TSS” peaks, all TAF4b “promoter-TSS” peaks for genes that were also DEGs, TAF4b “promoter-TSS” peaks for genes that were only Downregulated DEGs, and TAF4b “promoter-TSS” peaks for genes that were only Upregulated DEGs.

**Table S1. E14.5 RNA-seq analysis**.

**Table S2. E16.5 RNA-seq analysis**.

**Table S3. CUT&RUN data analysis**.

**Table S4. Cell numbers for experimental samples**.

